# Age-related DNA methylation changes are sex-specific: a comprehensive assessment

**DOI:** 10.1101/2020.01.15.905224

**Authors:** Igor Yusipov, Maria Giulia Bacalini, Alena Kalyakulina, Mikhail Krivonosov, Chiara Pirazzini, Noémie Gensous, Francesco Ravaioli, Maddalena Milazzo, Cristina Giuliani, Maria Vedunova, Giovanni Fiorito, Amedeo Gagliardi, Silvia Polidoro, Paolo Garagnani, Mikhail Ivanchenko, Claudio Franceschi

## Abstract

In humans, females live longer than males but experience a worse longevity, as genome-wide autosomal DNA methylation differences between males and females have been reported. So far, few studies have investigated if DNA methylation is differently affected by aging in males and females. We performed a meta-analysis of 4 large whole blood datasets, comparing 4 aspects of epigenetic age-dependent remodeling between the two sexes: differential methylation, variability, epimutations and entropy. We reported that a large fraction (43%) of sex-associated probes undergoes age-associated DNA methylation changes, and that a limited number of probes shows age-by-sex interaction. We experimentally validated 2 regions mapping in *FIGN* and *PRR4* genes, and showed sex-specific deviations of their methylation patterns in models of decelerated (centenarians) and accelerated (Down syndrome) aging. While we did not find sex differences in the age-associated increase in epimutations and in entropy, we showed that the number of probes showing age-related increase in methylation variability is 15 times higher in males compared to females. Our results can offer new epigenetic tools to study the interaction between aging and sex and can pave the way to the identification of molecular triggers of sex differences in longevity and age-related diseases prevalence.

## Introduction

A profound and multifaceted remodeling of DNA methylation patterns occurs during human aging [1–3]. DNA methylation profiles tend to diverge among individuals during life course [4–6], shaped by an intricate combination of environmental exposures, random events and genetically-driven mechanisms. At the same time, several epigenome-wide association studies (EWAS) have shown that a subset of the about 28 million CpG sites of the genome undergoes age-associated normative changes, i.e. reproducible hypermethylation or hypomethylation events that normally occur in all individuals during physiological, healthy aging (normative aging) [7,8]. Despite some controversial results [9,10], at least a fraction of normative epigenetic changes is tissue specific, indicating that the cellular microenvironment affects the activity of the molecular writers of DNA methylation patterns during aging. In the last 10 years an increasing number of studies identifyed age-associated DNA methylation changes at the level of single CpG sites, paving the way for the development of models, termed “epigenetic clocks”, that predict the age of an individual on the basis of their epigenetic profile [11]. Epigenetic clocks are an appealing resource for chronological age estimation in forensic applications, but they have risen to the limelight particularly because multiple reports have shown that they are sensitive to the health status of an individual and are thus informative of their biological age. Although a conclusive association between epigenetic clocks predictions and risk of age-related diseases is still missing [12], several independent studies showed that epigenetic age acceleration (i.e., predicted epigenetic age higher than effective chronological age) is associated to age-related diseases like cancer, cardiovascular disease and neurodegenerative conditions and to all-cause mortality [13]. On the other side, epigenetic age deceleration was reported to be associated with successful aging and longevity [14,15].

Surprisingly, the research on the DNA methylation changes occurring during aging has largely neglected one of the hot topics in aging research, i.e. the sex differences in lifespan and health span.

According to Global Health Observatory (GHO) data [16], global life expectancy at birth in 2016 was 74.2 years for females and 69.8 years for males and, although with different extent, this sex gap in longevity is worldwide [17]. At the same time, epidemiological data indicate that women live longer than men but experience a worse quality of life in advanced age [18]. Sex disparity exists for several diseases: cardiovascular disease, cancer and Parkinson’s disease have higher mortality rates in males than in females at a given age, while females are at higher risk of Alzheimer’s disease and show an increased prevalence of disabling conditions like bone and joint problems and autoimmune diseases. The reasons of these differences are still unclear, but it is likely that they result from a strict interplay between nature (for example, differences in hormones, asymmetries in genetic inheritance, sexual dimorphism) and nurture (for example, different vulnerability to environmental hazards, sexual selection) [19]. Notably, sex-specific longevity loci have been recently identified [20], further pointing out the contribution of sex on aging trajectories.

Independent studies reported DNA methylation differences between males and females in various tissues [21–23], involving CpG sites widespread across the autosomal chromosomes. These differences mirror the diversity in transcriptomic and proteomic profiles between the two sexes that have been recently reported [24,25]. However, few studies have investigated whether DNA methylation differences exist in whole blood between males and females during aging and whether they contribute to the sex gap in aging and longevity. Independent studies reported that according to Horvath’s clock males have an acceleration in epigenetic age compared to females [26–28]. Masser et al. analyzed genome-wide DNA methylation in mouse hippocampus and human frontal cortex and reported CpG sites that show different DNA methylation levels between males and females lifelong (referred as sex differences) and CpG sites that are differently affected by aging in males and females (referred as sex divergence) [29]. The vast majority of EWAS studies on aging have been performed in whole blood, but sex has usually been exiled as a confounding factor and used to adjust DNA methylation data.

In the present work we specifically investigated sex differences in whole blood DNA methylation changes during aging. We provide the results of a comprehensive study of four large whole blood datasets considering different aspects of age-associated epigenetic remodeling that can, either individually or in combination, contribute to the sex-specificity of human aging and longevity. In particular we focused on: 1) age-related changes in DNA methylation levels [8,30]; 2) age-related increase in DNA methylation variability, as described by Slieker et al. [6]; 3) age-related increase in epimutations, i.e. rare or stochastic changes in DNA methylation levels that are not shared among subjects, as defined by Gentilini et al. [5] 4) age-related increase in entropy in DNA methylation profiles, as previously described by Hannum et al. [31]. Furthermore, we investigated a subset of the loci emerged from these analyses in human models of successful and unsuccessful aging. Our results are compared with the only study that, to the best of our knowledge, recently investigated normative age-by-sex DNA methylation differences in whole blood of a single population, i.e. a large Scottish cohort [32].

## Results

### Identification of sex- and age-associated differentially methylated positions

We performed a meta-analysis of 4 large datasets on whole blood (Materials and Methods) to identify CpG sites with differential methylation between males and females (sex-associated differentially methylated positions, sDMPs). We identified 38100 sDMPs (Bonferroni corrected p-values resulting from meta-analysis <0.01), 53% of which were hypermethylated in females compared to males. We used the same datasets to identify age-associated differentially methylated positions (aDMPs) and we selected a list of 87581 probes (Bonferroni corrected p-values resulting from meta-analysis <0.01), 52% of which underwent hypermethylation with aging. We then asked how many sDMPs underwent DNA methylation changes with age, *i.e.* were also aDMPs. The intersection between sDMPs and aDMPs lists returned 16526 probes, that we defined sex- and age-associated differentially methylated positions (saDMPs); we defined the remaining probes (21574) sex-but *not* age-associated differentially methylated positions (snaDMPs). The proportion of sex-associated probes showing age-associated changes (16526 out 38100) was higher then expected, considering the proportion of age-associated probes in the genome (87581 out 327905; Fisher’s exact test p-value < 2*10-16, odds ratio 2.35). Figure 1A reports a graphical representation of the procedure used to identify saDMPs and snaDMPs. We found that most of the saDMPs undergoing hypomethylation with age were more methylated in males compared to females, while most of the age-hypermethylated saDMPs were more methylated in females compared to males (Figure 1B). Four examples of saDMPs are depicted in Figure 1C. The lists of saDMPs and of snaDMPs are reported in Supplementary File 1.

**Figure 1.**
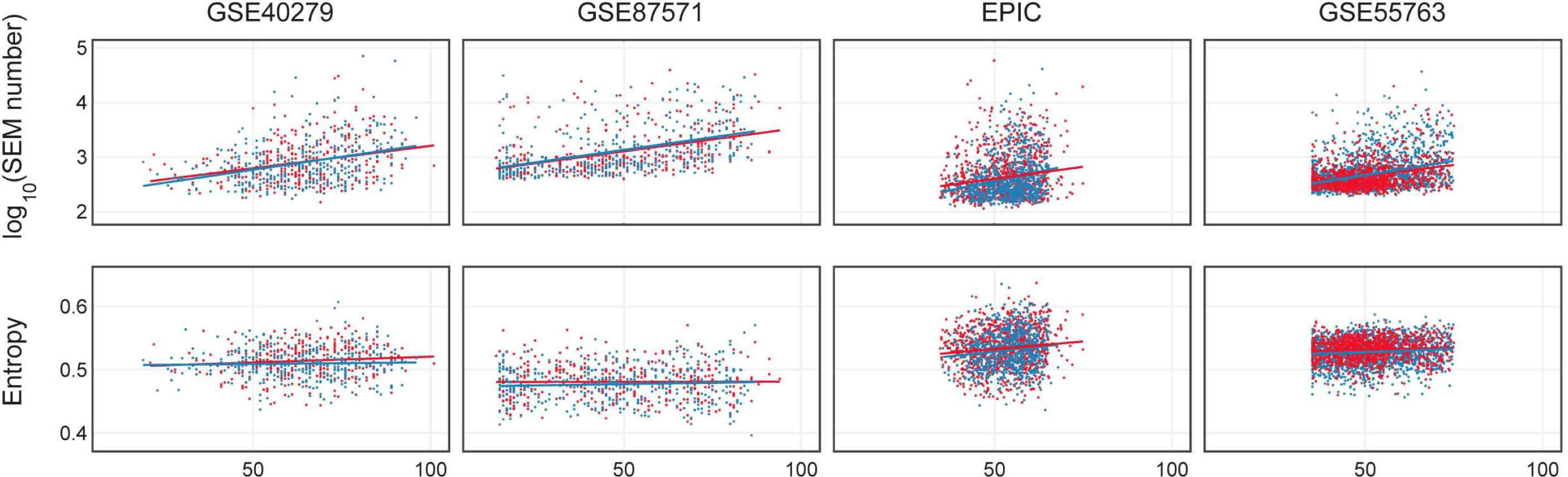
Identification of sex-specific age-associated differentially methylated positions (ssaDMPs). (A) The procedure used to identify saDMPs and snaDMPs. (B) Scheme of the number of sDMPs, aDMPs and saDMPs, divided according to the direction of methylation changes respect to sex (hyper- or hypo-methylated in males compared to females) and age (hyper- or hypo-methylated with increasing age). (C) Scatter plots of a selection of saDMPs: cg01620164 is hypomethylated in males and undergoes age-associated hypomethylation; cg03890691 is hypomethylated in males and undergoes age-associated hypermethylation; cg07128102 is hypermethylated in males and undergoes age-associated hypomethylation; cg04628369 is hypermethylated in males and undergoes age-associated hypermethylation.

When compared to previously published studies, we found that a total of 1121 saDMPs and 2163 snaDMPs were reported to have sex-dependent methylation (independently from age) in previous reports [21–23], also when newborns were considered [23] (Supplementary File 2).

We then investigated the possible functional role of saDMPs and snaDMPs. First of all, we explored whether the selected probes were enriched in specific genomic regions (Supplementary Figure 2), and we found that both saDMPs and snaDMPs were significantly enriched in Shore regions. The list of saDMPs, but not of snaDMPs, was significantly depleted in imprinted regions (p-value = 0.04, odds ratio: 0.64), as defined by Court et al. [33]. We also checked for an presence of sex hormone-related genes, as suggested by [22], in the lists of saDMPs and snaDMPs. saDMPs and snaDMPs mapped in 6610 genes and 8367 genes respectively. 2899 genes were shared between the two lists, indicating that the same gene can include multiple CpG sites with DNA methylation differences between males and females, only a subset of which shows also age-associated changes. The list of saDMPs included a higher proportion of hormone-related genes (28/6610 genes for saDMPs; 29/8367 for snaDMPs), but the enrichment was not statistically significant (p-value: 0.08; odds ratio: 1.6). Then, we analysed the two lists for their enrichment in gene ontologies according to GO database. While the list of snaDMPs was not enriched in any biological process (FDR corrected p-value<0.01), saDMPs were enriched in multiple ontologies related to neuronal and developmental functions and to cell-cell interactions (Supplementary File 3).

**Figure 2.**
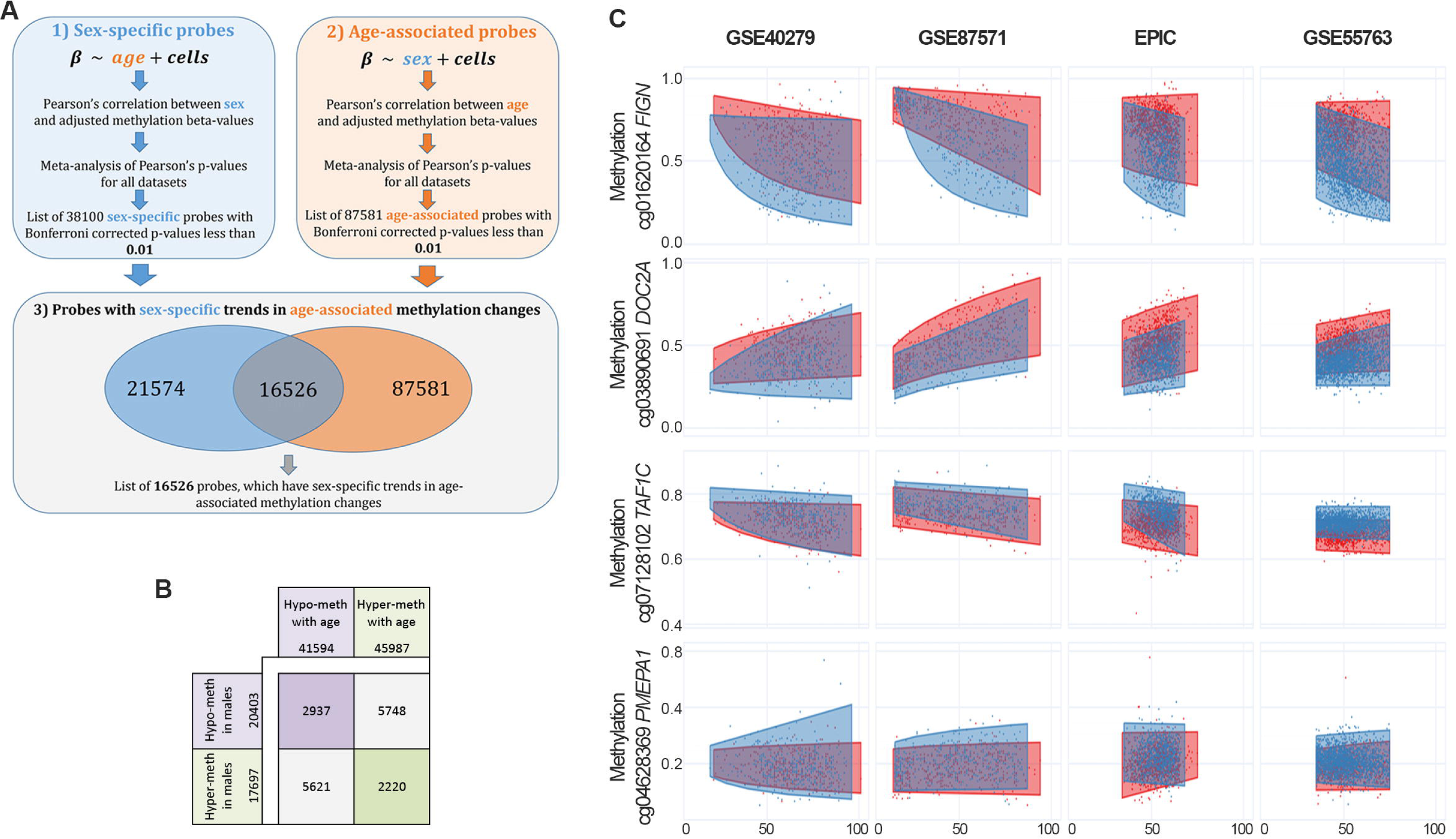
Validation of *FIGN* and *PRR4* loci by EpiTYPER. (A) Methylation of CpG unit 9 in *FIGN* amplicon *vs* age. (B) Methylation of CpG unit 3 in *PRR4* amplicon *vs* age. For each CpG unit, DNA methylation in controls (general population), centenarians, centenarian’s offspring and persons with Down syndrome is reported *vs* the age of the subjects. Males are in blue, females are in red. Linear regression between DNA methylation and age was calculated separately for males and females in control subjects and was reported in each plot.

Finally, we evaluated age-by-sex interactions in the 4 datasets. Meta-analysis resulted in 8 CpG probes whose methylation showed different aging trajectories according to sex (Supplementary File 4). Two of these CpG sites were previously identified as having age-by-sex interaction in whole blood [32], and the most significant CpG site (cg18834375) mapped within *FIGN* gene, that was the top ranker also in the list of saDMPs and included multiple CpG sites showing sex- and age-dependent methylation (cg01620164, cg19156483, cg10864319, cg18834375, cg15259986 and cg03878133).

### Validation of saDMPs

A subset of the above-identified saDMPs was experimentally validated using the EpiTYPER assay, a high throughput approach for target DNA methylation analysis. Target regions were chosen within *FIGN* and *PRR4*. We analyzed whole blood from 198 males from 15 to 98 years old and 221 females from 23 to 98 years old.

The *FIGN* target region included 13 CpG sites; of these, 7 were measurable by the assay, grouped in 5 CpG units. CpG unit 3.4.5 included the microarray probe cg01620164. We found that this group of CpG sites showed a sex-specific DNA hypomethylation trajectory comparable to the one resulting from the microarray (Supplementary Figure 3); also, the adjacent CpG sites showed a similar profile (Supplementary Figure 3), in particular CpG unit 9 (Figure 2A). This result indicates that the CpG sites located in this genomic region of at least 250bp are concordantly regulated during aging according to the sex of the individual in whole blood.

**Figure 3.**
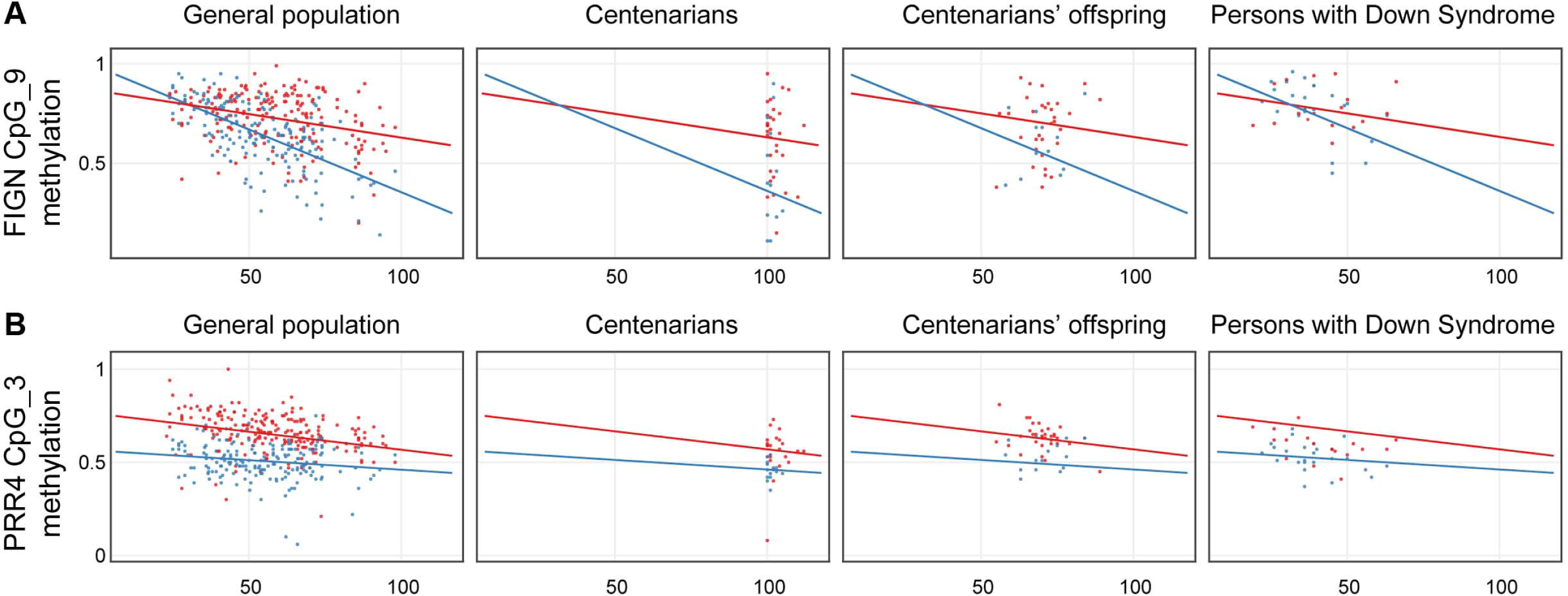
Identification of sex-specific age-associated variably methylated positions (ssaVMPs) (A) The approach used to indentify ssaVMPs. (B-E) Some examples of saVMPs showing age-associated increase in variability in males (B,C), age-associated decrease in variability in males (D) or age-associated increase in variability in females (E). *x* axis corresponds to age of subjects, *y* axis to methylation levels.

The *PRR4* target region included 5 CpG sites, all assessable by EpiTYPER and all corresponding to an Infinium450k probe; CpG units 3 and 4, corresponding to the Infinium450k probes cg23256579 and cg27615582, had the same mass and returned the same methylation value in the EpiTYPER assay. While CpG units 1 and 2 did not show age-dependent changes nor sex specificity (Supplementary Figure 4), CpG units 3 and 4 showed sex-dependent trajectories with aging (Figure 2B). Although less evident, also CpG 5 showed sex-related differences in age-associated methylation changes (Supplementary Figure 4).

**Figure 4.**
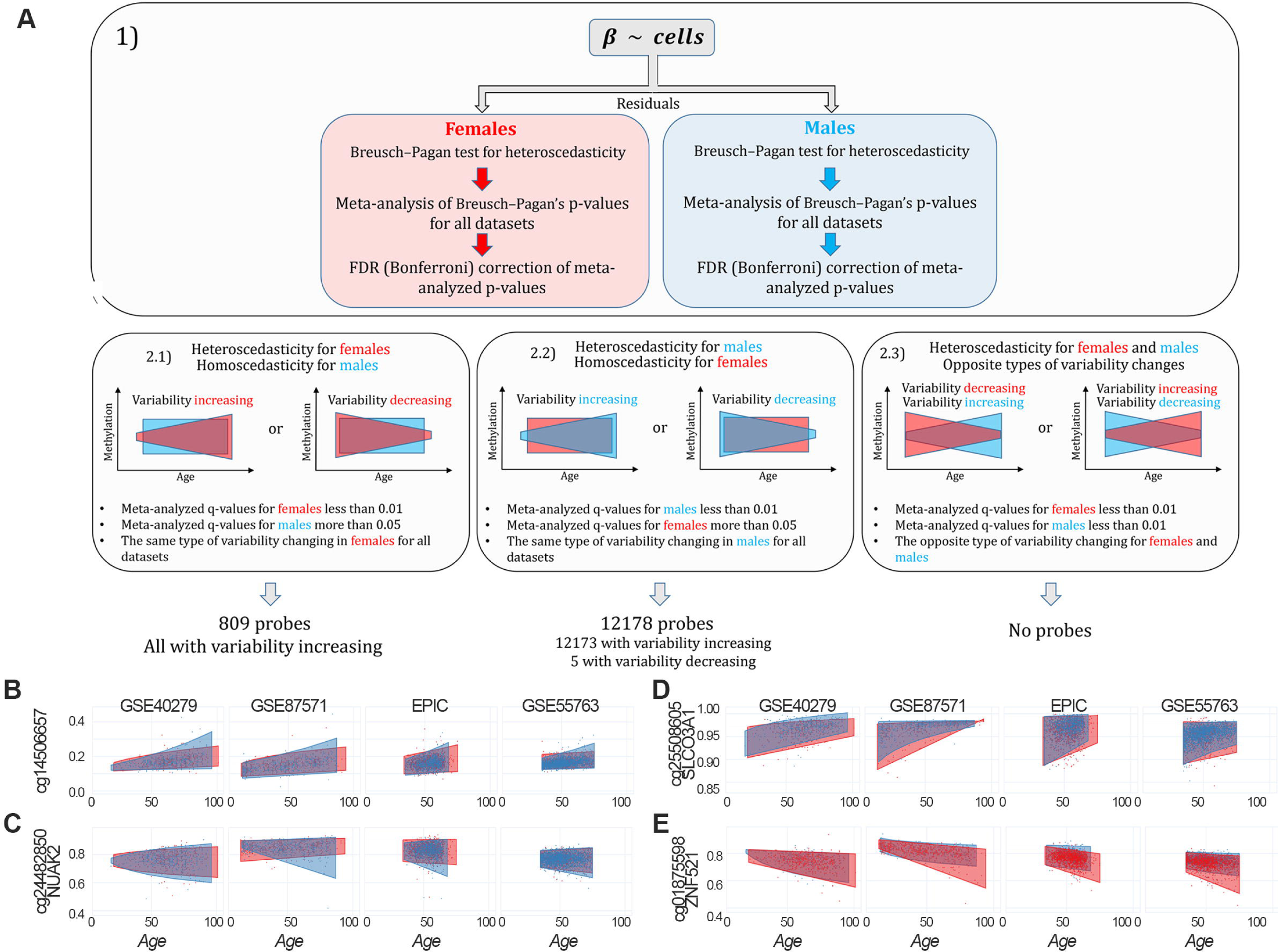
(A) Number of epimutations (log scale) in dependence on age in females (red) and males (blue). (B) Shannon entropy for 4 considered datasets: GSE40279, GSE87571, EPIC, GSE55763.

We used the EpiTYPER assay to investigate the two validated loci in samples from additional cohorts available in our laboratory: persons affected by Down syndrome, that we previously demonstrated to have an acceleration in epigenetic age [34,35]; and centenarians and their offspring, as a model of successful aging experiencing a deceleration in epigenetic age [14]. Interestingly, we found sex-dependent patterns of *FIGN* and *PRR4* methylation also in these models. Compared to aged controls (>80 years old), centenarian males displayed highly variable DNA methylation profiles for *FIGN* amplicon, with about half of the subjects showing a female-like DNA methylation level (Figure 2A and Supplementary Figure 5); the differences in variance between control and centenarians’ males (but not females) reached statistical significance for CpG unit 9 (F-test p-value: 0.02; Supplementary File 5). No specific trends were found for *PRR4* amplicon in centenarians’ cohort (Supplementary Figure 6 and Supplementary File 5). Centenarians’ offspring showed DNA methylation patterns comparable to age-matched controls for both the amplicons (Supplementary Figures 5 and 6). Persons affected by Down Syndrome showed DNA methylation profiles similar to age-matched controls in *FIGN* locus (Supplementary Figure 5). On the contrary, females affected by Down syndrome showed lower values of CpG unit 3 in *PRR4* amplicon compared to sex- and age-matched healthy controls (anova p-value correcting for age: 6.2*10-5; Supplementary File 5), while no significant differences were found between males affected by Down syndrome compared to sex- and age-matched controls (Figure 2B, Supplementary Figure 6 and Supplementary File 5). The results of the statistical analyses performed on the centenarians’, centenarians’ offspring and Down syndrome cohorts are summarized in Supplementary File 5.

### Identification of sex-specific age-associated variably methylated positions (ssaVMPs)

An increase in inter-individual DNA methylation variability has been described during aging, but a possible sex-specific effect has not been investigated. To have a general view of the sex-dependent trends in age-related increase in DNA methylation variability, we plotted the density distributions of standard deviation values, calculated in the GSE87571 dataset (the one with the most homogeneous distribution of ages) in 3 age-ranges (14-39 years; 40-59 years; 60-94 years) considering the whole cohort (Supplementary Figure 7A) or separating males and females (Supplementary Figure 7B). A clear increase in standard deviation across the 3 age ranges was evident when considering the entire cohort. No clear differences between males and females were evident in the first 2 age ranges, while we found a trend towards higher variability in males in the oldest group.

To identify probes having sex-specific differences in age-dependent variability (sex-specific age-associated variably methylated positions, saVMPs), we applied the approach described in Materials and Methods and reported in Figure 3A. We identified 809 and 12178 saVMPs specific for females and males respectively (Supplementary File 6). All the female-specific saVMPs displayed increased variability with age, and similarly only for 5 out 12178 male-specific saVMPs variability decreased with age. No probes with opposite trends in the two sexes were identified. Some examples of female- and male-specific saVMPs are reported in Figures 3B-E.

While female-specific saVMPs were enriched in Islands, male-specific saVMPs were enriched in Shore regions (Supplementary Figure 8). We also found that male-specific saVMPs were enriched in imprinted regions and mapped in 20 hormone-related genes, although this enrichment was not significant (p-value: 0.09; odds ratio: 1.66). Both female- and male-specific saVMPs were enriched in several gene ontologies related to neuronal and developmental processes, with some ontologies shared between the two lists (Supplementary file 7).

### Epimutations and entropy analysis

Epimutations were calculated in each dataset as previously described [5]. As shown in Figure 4A, we confirmed an increase in the number of epimutations with age both in males and females (p-value <0.01 in all the datasets), but no sex specific trends were found according to the second criterion of the “polygon” approach, described in Materials and Methods section.

The dependence of Shannon entropy on age for the 4 datasets is shown in Figure 4B. Entropy showed a significant increase with age (p-value <0.01) in males and females from the EPIC dataset and in females from the GSE87571 and GSE55763 datasets. Again, we applied the second criterion of “polygon” approach to entropy data, showing that there was no difference between sexes in Shannon entropy age-dependent increase.

## Discussion

Males and females experience different aging trajectories for several phenotypic traits [36,37]. Sex-specificity is established and maintained by differential genomic regulation, as evidenced by the profound transcriptomic, epigenomic and proteomic differences between males and females. However, how these differences in genome regulation evolve during life course has been poorly investigated, thus leaving a gap in our understanding of sexual dimorphism in aging and of its consequences in terms of morbidity and mortality.

In the present study we aimed at filling this gap by exploiting 4 large EWAS studies on human whole blood, including men and women of different ages and populations, in which we analyzed the sex specificity of age-associated normative changes, variability, epimutations and entropy.

The main findings we will discuss are the following: i) a large fraction of probes with sex-specific DNA methylation undergoes also hyper- or hypo-methylation during aging, and a small number of probes showed significant age-by-sex interaction; ii) the methylation of 2 selected saDMPs, aping in *FIGN* and *PRR4* genes, is differently modulated in centenarians and Down syndrome persons, assumed as human models of successful and unsuccessful aging [14,34,35]; iii) males display a higher number of saVMPs respect to females, the vast majority of which show age-associated increase in methylation variability; iv) males and females do not differ for the age-associated increase in epimutations and in entropy.

### i) saDMPs in healthy subjects of different ages and populations

We reported that 43% of probes showing sex-associated DNA methylation differences in whole blood display also age-associated changes (saDMPs). This result suggests that CpG sites showing DNA methylation differences between males and females are particularly prone to undergo epigenetic changes during aging. Interestingly we found that while saDMPs were enriched in gene ontologies related to neuronal and developmental functions, sex-but not age-associated probes were not enriched in any particular biological process. It is difficult to speculate the mechanisms leading to differential methylation of these genes in males and females and to age-associated changes. When considering autosomal differences of DNA methylation between men and women (correcting for age), Singmann et al. found an enrichment in CpG island shores and in imprinted genes, but they did not find an enrichment in sex hormone-related genes [22]. Our list of saDMPs was enriched in CpG island shores but depleted in imprinted genes. We did not find any significant enrichment in sex-hormone related genes, although saDMPs included a higher proportion of sex-hormone related genes compared to snaDMPs.

Finally, we searched for those probes displaying different DNA methylation trajectories in males and females during aging. We identified 8 probes having age-by-sex interaction, 2 of which were recently reported by McCartney et al in an independent dataset [32]. In particular we found that *FIGN* gene, which encodes for an ATP-dependent microtubule severing enzyme which catalyzes internal breaks in microtubules, included both probes from the saDMPs list and displaying age-by-sex interactions.

Future studies should assess whether the DNA methylation patterns that we described are modulated by the changes in sex hormones during life-course and if the different chromosomal repertoire of males and females can affect age-related epigenetic trajectories and/or the effect of environment on epigenetic profiles.

### ii) saDMPs in centenarians and Down syndrome persons

Another question is if and how the saDMPs that we identified contribute to the sex gap in health span and longevity. To this aim, we exploited two cohorts available in our lab, in which we measured whole blood DNA methylation by the targeted EpiTYPER assay: persons affected by Down syndrome, as a model of premature/accelerated aging [34,35], and centenarians, as a model of successful/decelerated aging [14]. The results are intriguing, as both models showed a peculiar sex specific alteration in *FIGN* and *PRR4* epigenetic patterns. In particular, a subset of centenarian males showed a “feminization” of *FIGN* methylation values, while females with Down syndrome showed a “masculinization” of *PRR4* methylation values. No differences were found in the centenarians’ offspring group, despite we and others previously showed an epigenetic age deceleration effect in these subjects [14]. Interestingly, the “feminization” of centenarians methylation patterns at *FIGN* locus is reminiscent of the gene expression shift towards female patterns observed after caloric restriction [38,39]. Further studies should deepen these results and identify other changes in saDMPs that are associated to age-related diseases or longevity.

### iii) saVMPs in healthy subjects of different ages and populations

An increase in epigenetic variability has been reported during aging [6], in line with what observed for other molecular layers [40–42]. In the GSE87571 cohort we showed a global increase in DNA methylation variance during aging, and we further reported a trend towards higher variance in males compared to females at older ages. This result mimics what observed for gene expression in the hippocampus of male and female mice at different ages [43], thus suggesting that the loss of epigenetic and transcriptional control that occurs during aging is more marked in males than in females. Accordingly, a more specific search for saVMPs showed that in males the number of probes showing age-related changes in methylation variability is 15 times higher than in females and that the vast majority of these probes displays and increase in age-realted variability, as previously reported [6]. Interestingly, this list was significantly enriched in imprinted regions. An increase in variability at these loci can be related to the phenomenon of loss of imprinting, which has been largely reported in cancer and demonstrated to occur during aging [44–46].

### iv) Epimutations and entropy

While variable probes are defined at the level of population, epimutations are rare methylation changes that are specific for one or few individuals within a certain population. As such, variable probes and epimutations represent distinct aspects of epigenetic instability, that can be differently triggered during aging and that can differently affect aging trajectories. Accumulation of epimutations has been reported in cancer [47], and we and others demonstrated that number of epimutations increases with age [5,48]. Recently, Wang and colleagues showed that the number of epimutations in whole blood tends to be higher in females compared to males [48]. On the contrary, we failed to detect differences in the age-related increase in epimutations between the two sexes. This discrepancy can be due to the different analytical approaches and/or to cohort-specific effects, as Wang et al investigated monozygotic and dizygotic twins longitudinally assessed. A recent paper assessed epimutations in 3 large cohorts and did not find significant differences between males and females [49]. Similarly, we did not find sex-related differences in age-related changes in Shannon entropy, another measure of epigenetic drift.

### Strengths and Limitations

The main strengths of our work are: i) we compared the two sexes not only for age-associated hyper- or hypo-methylation changes, but also for other types of epigenetic remodeling (variability, epimutations, entropy) that, although less characterized, are likely to affect aging and be involved in the sex gap in longevity; ii) the analysis was performed independently in 4 distinct datasets including subjects recruited in different geographic area (United States, Sweden, Italy, United Kingdom) and belonging to different ancestries (European and Hispanic); this contributes to disentangle the effects of sex from those of potentially confounding factors like genetic background and socio-cultural aspects related to gender definition; iii) beside healthy individuals representative of physiological aging, we evaluated also extreme phenotypes (persons with Down Syndrome, centenarians and their offspring) that provide a first descriptive insight on the possible contribution of sex-specific methylation in the sex gap in aging and longevity.

At the same time, our study has some limitations. The analyzed datasets differ in terms of size, age-range and data pre-processing procedures (in particular the GSE40279 dataset). It is therefore likely that our selection excluded additional CpG sites displaying a sex-specificity in their age-associated methylation trends, but not evident in all the datasets due to the above-mentioned differences between them. Furthermore, although some ancestries are included, many are missing, and the study of additional populations will be necessary to distinguish the effects of sex and gender in shaping age-related methylation changes. Finally, similarly to [22], our study includes only autosomal probes (after the removal of cross-reactive probes), while recently McCartney et al found that many of the probes with a sex by age interaction are on the X chromosome [32].

### Conclusions and future perspectives

In conclusion, we provided a comprehensive description of sex-differences in DNA methylation changes with aging in whole blood. Our results suggest that a large fraction of CpG sites with sex-specific DNA methylation patterns are also modulated during aging, and that sex can affect some aspects of age-related epigenetic remodeling, like increase in variability in DNA methylation patterns. Future studies should investigate the tissue-specificity of these patterns and their relationship with gene expression differences between males and females, in particular for those probes that show age-by-sex interactions, in order to identify possible molecular triggers of sex gap in aging and longevity. Importantly, here we reported also a list of sex-but not age-associated probes, and we cannot exclude that also these sites can contribute to the dimorphism in aging phenotypes between males and females.

Our results pave the way for the development of a new generation of sex-specific epigenetic clocks that, comparing to the “unisex” clocks currently available, are likely to be more informative of the peculiar trajectories that males and females experience during aging. Thus, these epigenetic tools will be valuable in the study of age-related diseases with different prevalence in males and females (cancers, cardiovascular, autoimmune and neurodegenerative diseases) and of those conditions (intersex, hormone therapies, transgender, etc) in which the interaction between sex/gender and aging has not yet been adequately investigated.

## Materials and Methods

### Datasets

The Gene Expression Omnibus (GEO) Datasets repository [50] was interrogated using “GPL13534” (the accession code of the platform HumanMethylation450 BeadChip, Illumina) and “blood” as search terms, setting “tissue”, “age”, “gender” and “sex” as attributes and sorting the results by Number of Samples (High to Low). Only datasets including healthy subjects were considered. Based on these criteria, as to June 1st 2019 we selected the 3 datasets including the highest number of samples: GSE40279 [31], GSE87571 [51] and GSE55763 [52]. Furthermore, we analyzed a fourth dataset not uploaded in GEO that is part of the EPIC Italy study [53]. The total number of subjects included in each dataset, as well as the number of males and females, is reported in Supplementary Table 1. Supplementary Figure 1 reports, for each dataset, the number of males and females according to age.

For EPIC dataset, raw data were normalized using an in-house software written for the R environment and extensively described in [54]. For the datasets downloaded from GEO, raw data (.idat files) were available only for GSE87571. We extracted these .idat files using *minfi* Bioconductor package and normalized them using the *preprocessFunnorm* function implemented in the same package [55]. For the remaining datasets, the analyses were performed on pre-processed beta value matrixes available in GEO: according to authors’ indications, the GSE55763 data were the result of a quantile normalization of intensity values, while GSE40279 beta values were not normalized but adjusted for internal controls by the Illumina’s Genome Studio software.

Probes mapping on sex chromosomes and probes with internal SNPs, with non-unique mapping to the bisulfite-converted genome and with off-target hybridization according to [56] were excluded from each dataset, leaving 414505 probes for GSE40279, 414950 probes for GSE87571, 349534 probes for EPIC and 382458 probes for GSE55763. 327905 probes were common to the four datasets and were considered in the analyses described below.

In each dataset, blood cell proportions were estimated from methylation data using Horvath calculator [57].

### Identification of age-associated probes having sex-specific DNA methylation patterns

To identify CpG sites showing DNA methylation differences between the two sexes *and* age-associated changes in DNA methylation (sex- and age-associated differentially methylated positions, saDMPs), we proceeded as follows (Figure 1A): 1) We meta-analyzed the 4 datasets to identify probes with sex-dependent DNA methylation patterns (sDMPs). As previously described [22], in each dataset we calculated the p-value of Pearson’s correlation between sex and methylation beta-values, previously adjusted for age and blood cell proportions (CD8T cells, CD4T cells, NK cells, B cells and granulocytes, estimated as described above). METAL [58] was used to perform sample-size weighted meta-analysis on the 4 lists of Benjamini-Hochberg corrected p-values. 2) We meta-analyzed the 4 datasets to identify probes with age-dependent DNA methylation patterns (aDMPs). In each dataset, we calculated the p-value of Pearson’s correlation between age and methylation beta-values, previously adjusted for sex and blood cell proportions (CD8T cells, CD4T cells, NK cells, B cells and granulocytes, estimated as described above). As described above, METAL [58] was used to perform sample-size weighted meta-analysis on the 4 lists of Benjamini-Hochberg p-values. 3) The meta-analyzed p-values were corrected using Bonferroni procedure and a significance threshold of 0.01 was considered. Furthermore, only probes with a concordant trend in all the 4 datasets (for sDMPs: hypermethylated or hypomethylated in males respect to females in all the datasets; for aDMPs: hypermethylated of hypomethylated with age in all the datasets) were considered, returning a list of 38100 sDMPs and a list of 87581 aDMPs. 4) Finally, to identify saDMPs, we intersected the list of sDMPs and the list of aDMPs, resulting in a list of 16526 probes.

### Identification of probes having sex-specific trends in age-associated methylation variability

To identify probes having sex-specific differences in age-dependent variability of methylation (sex-specific age-associated variably methylated positions, ssaVMPs) we proceeded as follows (Figure 3B): 1) We first regressed out the estimates of CD8T cells, CD4T cells, NK cells, B cells and granulocytes from beta values in each dataset; 2) in order to check for heteroscedasticity respect to age, we applied the Breusch-Pagan for males and females separately in each dataset [59,60]. 3) Heteroscedasticity p-values were analyzed with sample-size weighted meta-analysis using METAL [58] for males and females separately. The meta-analyzed p-values were corrected using Bonferroni procedure and a significance threshold of 0.01 was considered. 4) We defined 3 possible scenarios of sex-specific differences in age-dependent variability of methylation: a) Probes heteroscedastic in females and homoscedastic in males, that is probes with a meta-analyzed and Bonferroni-corrected p-value less than 0.01 in females and higher than 0.05 in males. b) Probes homoscedastic in females and heteroscedastic in males, that is probes with a meta-analyzed and Bonferroni-corrected p-value less than 0.01 in males and higher than 0.05 in females. c) Probes heteroscedastic in both females and males, but with opposite directions of change in variability (variability increases in females and decreases in males or variability decreases in females and increases in males).

### Identification of epimutations and Shannon entropy analysis

To identify epimutations (*i.e.*, CpG probes for which one or few individuals show extremely different methylation levels compared to the rest of the cohort), for each probe we calculated the interquartile ranges of beta values; we then selected the probes having one or more subjects (epimutated subjects) having a beta value exceeding three times interquartile ranges (Q1 – (3×IQR) and Q3 + (3×IQR)), as reported in [5].

To calculate Shannon entropy, we applied the following procedure, according to [61]: 1) we obtained residuals by filtering out the dependence of beta-values on age and blood cells proportions; 2) we recalculated beta-values according to the formula:

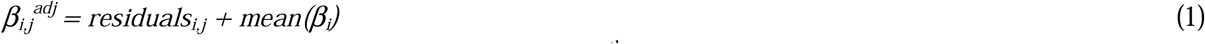

where *mean(β_*i*_)* is the average methylation level for i^th^ CpG site, *i* is the index of CpG and j is the index of subject. Then, we calculated Shannon entropy using the following formula, as indicated in [31]:

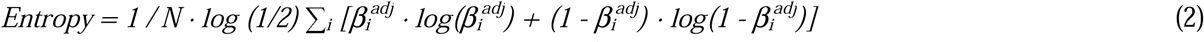

where 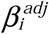 is the recalculated methylation level for i^th^ CpG site and *N* is the number of CpG sites.

### Gene-targeted DNA methylation analysis

The EpiTYPER assay (Agena) was used to measure DNA methylation of *FIGN* and *PRR4* in whole blood from 560 subjects belonging to 4 groups: 419 healthy controls of different ages, 49 centenarians, 48 centenarians’ offspring and 44 persons with Down Syndrome. Age range and sex distribution of the 4 cohorts are reported in Supplementary Table 2. All the subjects were recruited following the approval by the Ethical Committee of Sant’Orsola-Malpighi University Hospital (Bologna, Italy).

Genomic DNA was extracted using the QIAamp 96 DNA Blood Kit (Qiagen) and 500 ng were bisulphite converted using EZ-96 DNA Methylation Kit (Zymo Research Corporation). Ten ng of bisulphite-converted DNA were amplified using the following bisulphite-specific primers, containing tag sequences for the EpiTYPER protocol: *FIGN forward* aggaagagagTTTTTTGAAAAGAGAGAAAGAAGGA; *FIGN reverse* cagtaatacgactcactatagggagaaggctATAAACAATCAAACCATCCAATTTCTA; *PRR4 forward* aggaagagagTTTGTGTTTTGAGTTGAGTTTAGAG; *PRR4 reverse* cagtaatacgactcactatagggagaaggctCCTAAAAATAAAACTTCTATCATCCA. Primers for *FIGN* and *PRR4* amplified chr2:164,589,883-164,590,418 and chr12:11,001,978-11,002,636 (GRCh37/hg19 genome assembly) respectively.

### Enrichment analyses

Enrichment of genomic regions, imprinted genes and sex-hormone related genes was calculated using Fisher exact test, as implemented in the *fisher.test* function in the *stats* R package. The lists of imprinted genomic regions [33] and of sex hormone-related genes used as background are reported in Supplementary File 1. Enrichment of GO annotations was calculated using the *methylgometh* function implemented in the *methylGSA* R package, using default settings [62].

## Abbreviations

sDMPs: sex-associated differentially methylated positions
aDMPs: age-associated differentially methylated positions
saDMPs: sex- and age-associated differentially methylated positions
snaDMPs: sex-but *not* age-associated differentially methylated positions
saVMPs: sex-specific age-associated variably methylated positions

## Author’s contributions

IY, MGB, MI, and CF contributed to the conception and design of the study. IY, CP, CG, GF, AG, SP organized the datasets. IY, MGB, AK, MK, MV, CP, CG performed the statistical analysis. MGB, NG, FR, MM, CP, CG performed the validation experiments. IY, MGB, PG, MI and CF wrote the manuscript. All authors contributed to manuscript revision, and read and approved the submitted version.

## Supplementary tables

**Supplementary Table 1.** Characteristics of the Infinium450k datasets investigated in the present study

|  | GSE40279 | GSE87571 | EPIC | GSE55763 |
| --- | --- | --- | --- | --- |
| Number of subjects | 656 | 729 | 1803 | 2670 |
| Number of females | 338 | 388 | 1114 | 860 |
| Number of males | 318 | 341 | 689 | 1810 |
| Age range | 19 – 101 | 14 – 94 | 34 – 74 | 35 – 75 |

**Supplementary Table 2.** Characteristics of the samples analysed by the EpiTYPER assay

|  | Healthy controls<br>n=419 | Centenarians<br>n=49 | Centenarians' offspring<br>n=48 | Down syndrome<br>n=44 |
| --- | --- | --- | --- | --- |
| Males | n=198<br>25-98 years | n=15<br>100-105 years | n=15<br>58-84 years | n=25<br>22-63 years |
| Females | n=221<br>23-98 years | n=34<br>100-112 years | n=33<br>55-89 years | n=19<br>19-66 years |

## Supplementary Figures

Supplementary Figure 1. Histograms of the number of females (red) and males (blue) according to age in GSE40279, GSE87571, EPIC and GSE55763 datasets.

Supplementary Figure 2. Enrichment (odds ratio) of genomic localizations for saDMPs (A) and snaDMPs (B).

Supplementary Figure 3. Validation of *FIGN* locus by EpiTYPER. For each of the CpG units returned by the EpiTYPER assay, DNA methylation in controls (general population), centenarians, centenarian’s offspring and persons with Down syndrome is reported *vs* the age of the subjects. Males are in blue, females are in red. Linear regression between DNA methylation and age was calculated separately for males and females in control subjects and was reported in each plot.

Supplementary Figure 4. Validation of *PRR4* locus by EpiTYPER. For each of the 5 CpG units returned by the EpiTYPER assay, DNA methylation in controls (general population), centenarians, centenarian’s offspring and persons with Down syndrome is reported *vs* the age of the subjects. Males are in blue, females are in red. Linear regression between DNA methylation and age was calculated separately for males and females in control subjects and was reported in each plot.

Supplementary Figure 5. Boxplots of DNA methylation for each CpG unit in *FIGN* amplicon in centenarians, centenarians’ offspring and Down syndrome cohorts. Left panels: for each CpG unit in *FIGN* locus, boxplots of DNA methylation in male and female centenarians, compared to male and female controls (>80, < 100 years). Middle panels: for each CpG unit in *FIGN* locus, boxplots of DNA methylation in male and females centenarians’ offspring, compared to age-matched male and female controls (>54, < 90 years). Right panels: for each CpG unit in *FIGN* locus, boxplots of DNA methylation in male and female Down syndrome persons, compared to age-matched male and female controls (>18, < 67 years).

Supplementary Figure 6. Boxplots of DNA methylation for each CpG unit in *PRR4* amplicon in centenarians, centenarians’ offspring and Down syndrome cohorts. Left panels: for each CpG unit in *PRR4* locus, boxplots of DNA methylation in male and female centenarians, compared to male and female controls (>80, < 100 years). Middle panels: for each CpG unit in *PRR4* locus, boxplots of DNA methylation in male and females centenarians’ offspring, compared to age-matched male and female controls (>54, < 90 years). Right panels: for each CpG unit in *PRR4* locus, boxplots of DNA methylation in male and female Down syndrome persons, compared to age-matched male and female controls (>18, < 67 years).

Supplementary Figure 7. Density distribution of standard deviation values calculated in the GSE87571 dataset for 3 age classes, considering males and females together (A) or separated (B).

Supplementary Figure 8. Enrichment (odds ratio) of genomic localizations for ssaVMPs calculated from beta values (A) or residuals (B).

